# Non-Communicable Diseases and their determinants: A Cross-sectional State-Wide STEPS Survey, Haryana, North India

**DOI:** 10.1101/482117

**Authors:** JS Thakur, Gursimer Jeet, Ria Nangia, Divya Singh, Sandeep Grover, Tanica Lyngdoh, Arnab Pal, Ramesh Verma, Ramnika Aggarwal, Mohd. Haroon Khan, Rajiv Saran, Sanjay Jain, KL Gupta, Vivek Kumar

**Affiliations:** Department of Community Medicine, Post Graduate Institute of Medical Education and Research, Chandigarh, India; Department of Psychiatry, Post Graduate of Medical Education and Research, Chandigarh, India; Indian Institute of Public Health Association, Public Health Foundation of India, Gurugram, India; Department of Biochemistry, Post Graduate Institute of Medical Education and Research, Chandigarh, India; Department of Social and Preventive Medicine, Post Graduate Institute of Medical Sciences, Rohtak, India; Department of Community Medicine, Kalpana Chawla Medical College, Karnal, India; Department of Community Medicine, Shaheed Hasan Khan Mewati Government Medical College, Mewat, India; Department of Internal Medicine and Epidemiology, University of Michigan, Ann Arbor, Michigan, USA; Department of Internal Medicine, Post Graduate Institute of Medical Education and Research, Chandigarh, India; Department of Nephrology, Post Graduate of Medical Education and Research, Chandigarh, India

**Author notes:** Corresponding author Dr. JS Thakur Professor of Community Medicine, Department of Community Medicine and School of Public Health, Post Graduate Institute of Medical Education and Research, Chandigarh, India Email Id Contributions: Conceptualization, Data curation, Formal analysis, Funding acquisition, Investigation, Methodology, Project administration, Resources, Software, Supervision, Validation, Visualization, Writing – original draft, Writing – review & editing. **Contact Addresses of all authors**: Dr. Gursimer Jeet Department of Community Medicine and School of Public Health, Post Graduate Institute of Medical Education and Research, Chandigarh, India Contributions: Conceptualization, Data curation, Formal analysis, Investigation, Methodology, Project administration, Resources, Software, Supervision, Validation, Visualization, Writing – original draft, Writing – review & editing; Ms. Ria Nangia Department of Community Medicine and School of Public Health, Post Graduate Institute of Medical Education and Research, Chandigarh, India Contributions: Data Curation, Formal analysis, Methodology, Project administration, Software, Supervision, Validation, Visualization, writing – original draft, Writing – review & editing; Ms. Divya Singh Department of Community Medicine and School of Public Health, Post Graduate Institute of Medical Education and Research, Chandigarh, India Contributions: Formal analysis, Methodology, Project administration, Resources, Software, Visualization, Writing – original draft; Dr. Sandeep Grover Department of Psychiatry, Post Graduate of Medical Education and Research, Chandigarh, India Contributions: Formal analysis, Methodology, Project administration, Resources, Software, Validation, Visualization, Writing – original draft, Writing – review & editing; Dr. Tanica Lyngdoh Indian Institute of Public Health Association, Public Health Foundation of India, Gurugram, India Contributions: Formal analysis, Investigation, Methodology, Project administration, Resources, Supervision, Validation, Visualization, Writing – review & editing; Dr. Arnab Pal Department of Biochemistry, Post Graduate Institute of Medical Education and Research, Chandigarh, India Contributions: Formal analysis, Investigation, Methodology, Resources, Software, Supervision, Visualization, Writing – review & editing; Dr. Ramesh Verma Department of Social and Preventive Medicine, Post Graduate Institute of Medical Sciences, Rohtak, India Contributions: Formal analysis, Investigation, Methodology, Project administration, Resources, Software, Supervision, Visualization, Writing – review & editing; Dr. Ramnika Aggarwal Department of Community Medicine, Kalpana Chawla Medical College, Karnal, India Contributions: Investigation, Methodology, Project administration, Software, Supervision, Validation, Visualization, Writing – review & editing; Dr. Mohd. Haroon Khan Department of Community Medicine, Shaheed Hasan Khan Mewati Government Medical College, Mewat, India Contributions: Investigation, Methodology, Project administration, Resources, Supervision, Validation, Writing – review & editing; Dr. Rajiv Saran Department of Internal Medicine and Epidemiology, University of Michigan, Ann Arbor, Michigan, USA Contributions: Formal analysis, Investigation, Project administration, Resources, Supervision, Visualization, Writing – review & editing; Dr. Sanjay Jain Department of Internal Medicine, Post Graduate Institute of Medical Education and Research, Chandigarh, India Contributions: Investigation, Methodology, Project administration, Software, Supervision, Validation, Visualization, Writing – review & editing; Dr. KL Gupta Department of Nephrology, Post Graduate of Medical Education and Research, Chandigarh, India Contributions: Methodology, Project administration, Resources, Supervision, Validation, Visualization, Writing – review & editing; Dr. Vivek Kumar Department of Nephrology, Post Graduate of Medical Education and Research, Chandigarh, India Contributions: Project administration, Resources, Supervision, Validation, Visualization, Writing – review & editing.

**Keywords:** Non-communicable Diseases, chronic diseases, risk factor survey, surveillance, STEPS, population based survey, cross-sectional survey, chronic disease, cross-sectional survey

## Abstract

**Background:** Recent studies have documented high variation in epidemiologic transition levels among Indian states with noncommunicable disease epidemic rising swiftly. However, the estimates suffer from non-availability of reliable data for NCDs from sub populations. In order to fill the knowledge gap, the distribution and determinants of NCD risk factors were studied along with awareness, treatment and control of NCDs among the adult population in Haryana, India.

**Methods:** NCD risk factors survey was conducted among 5078 residents, aged 18-69 years during 2016-17. Behavioural risk factors were assessed using STEPS instrument, administered through an android software (mSTEPS). This was followed by physical measurements using standard protocols. Finally, biological risk factors were determined through the analysis of serum and urine samples.

**Results:** Males were found to be consuming tobacco and alcohol at higher rates of 38.9% (95% CI: 35.3-42.4) and 18.8% (95% CI: 15.8-21.8). One-tenth (11%) (95% CI: 8.6-13.4) of the respondents did not meet the specified WHO recommendations for physical activity for health. Around 35.2% (95%CI: 32.6-37.7) were overweight or obese. Hypertension and diabetes were prevalent at 26.2% (95% CI: 24.6-27.8) and 15.5% (95% CI: 11.0-20.0). 91.3% (95% CI: 89.3-93.3) of the population had higher salt intake than recommended 5gms per day.

**Conclusion:** The documentation of strikingly high and uniform distribution of different NCDs and their risk factors in state warrants urgent need for evidence based interventions and advocacy of policy measures.

## Introduction

Consequent to world-wide declaration of war against NCDs [1], a national program on Noncommunicable diseases (NCDs), [2] was introduced in India by the end of last decade. Being a diverse country having 17% of world’s population [3], the pattern, distribution of diseases and their determinants vary a lot, ipso facto effecting the choice and delivery of evidence based prevention and control interventions. Being a country of diversities, one size fits all principle never fits for implementation of interventions in Indian states. In addition, lack of reliable estimates of distribution of risk factors warrant careful introspection into the landscape of epidemic in the different populations. Subsequent to the adoption of national monitoring framework [1], collection of data points for monitoring NCDs and their risk factors gained momentum in India. [4] Health system reforms for adapting exiting and establish new mechanisms for gearing up efforts towards the battle against NCDs still needs a push as baseline estimates of risk factors level in populations across different states remains unknown for many of the states. Interestingly, a report by NITI Aayog, a prominent national institution for policy formulation in India lamented upon the lack of availability of acceptable quality data to address critical areas such as, NCDs, mental health, governance, and financial risk protection in a health index for its states.[5] Another report highlights the need for local data for robust sub national estimates.[6] In order to fill the knowledge gap in a large Indian state, NCD risk factors survey was undertaken. The overall aim of the survey was to generate state specific NCD risk factors related information for use by program managers, explore the key determinants of NCD risk factors and policy makers along with capacity development within state to undertake these surveys in future. The World Health Organization (WHO)STEP wise approach to Surveillance (STEPS) of risk factors [7] was used to conduct the survey with the objective to determine the prevalence and distribution of the NCD risk factors in Haryana for 18-69 years old adult population. Collaborative arrangements were made to strengthen state’s capacity.

Haryana, a North Indian state houses 25 million Indians which constitutes 2% of nation’s population. The population number is comparable to many countries in developing as well as developed worlds such as Australia, Netherlands, Greece, Sri Lanka, Nepal, and Syria. This marks the importance of sub national surveys in India.[8] As per a recent report, the epidemiological transition level (ETL) of Haryana is 0.4.[6] Among different Indian states, it currently has the higher-middle ETL. Consequently, the state is currently faced with several socio economic development challenges including health. Despite documentation of high burden by some researchers, state wide surveys for non-communicable diseases could never be conducted. The systematic inputs into the morbidity burden due to NCDs, therefore, lies unknown for the state till date.[6]

## Methods

The state wide survey was a collaborative effort between five public and private funded medical and research institutes with Post Graduate Institute of Medical Education and Research, Chandigarh (PGIMER) as the planning and executing agency. The study was carried out among adults 18-69 aged years residing in Haryana State, India. The total duration of the study was 15 months out of which the population was surveyed for 5 months. The survey was designed in accordance with the WHO STEPwise approach to surveillance of NCDs (STEPS) [9] to provide prevalence estimates of risk factors for two age groups (18-44, 45-69 years) by gender and place of residence (urban/rural). A multistage, geographically clustered, probability-based sampling approach was used. The ultimate sampling units were the households and one individual residing in the selected household was selected using the Kish method.[10]

Using population estimates for each age group by gender cluster for the combined population of Haryana (based on the 2011 population census), sample size estimates were calculated for each age/gender strata. Since multistage cluster sampling method was used, the design effect for the survey was taken as 1.5 as recommended.[9] The total minimum sample size estimate (obtained by summing across the age/gender strata) of 5122 was obtained after adjusting for the design effect and for the expected response rate at 90%. Choice of very low expected non response rate was based on the previous experience of survey implementers in an adjacent state where response rate of 95% was achieved through adoption of innovative measures. Keeping minimum required sample size and number of Primary Sampling Units (PSUs) to be covered (150) in mind, 35 respondents per primary sampling unit were selected. Thus making a total pool of 5250 respondents across 150 units in Haryana.A total of 1607 wards and 6642 villages were included in the sampling frame. A proportionate allocation as per census distribution in urban and rural areas were adopted. In urban areas, a three-stage procedure was followed. In the first stage, wards were selected with probability proportional to size (PPS). In the second stage, one Census Enumeration Block (CEB) was randomly selected from each sampled ward. In the final stage households were randomly selected within each CEB using the systematic random sampling procedure. From each selected PSUs, i.e. village in rural area and CEB in urban area 35 households were selected. From each selected household, one individual was selected from those who fall in the 18-69 age range by using Kish method. (Supplementary file 2) Step 3 was conducted on half of sub-sample considering resource constraints.

The questionnaire for the survey was developed with adaptation of WHO STEPwise Surveillance (STEPS) version 3.1 questionnaire. Translation to Hindi and back translation in English was done. An android based application software (mSTEPS), piloted earlier was adapted for this survey.[11] Data collected for all steps was entered using the application. Socio demographic and behavioural information was collected in Step 1. Physical measurements such as height, weight, blood pressure, skinfold thickness, hip and waist circumference was collected in Step 2 using standardized instruments and protocols. Biochemical measurements were conducted on serum and urine samples to assess fasting blood glucose, total cholesterol, triglycerides, serum creatinine and albumin-creatinine ratio in Step 3 using a mix of wet and dry chemistry methods.

### Data Collection (STEP 1)

All the selected field investigators/interviewers underwent a four-day training for collecting data of all the three Steps before the survey. The training was imparted on 11 domains using the mSTEPS application and included interactive sessions, discussions and hands-on training for physical measurements. Methodology of selecting, notifying and approaching the household/ respondent was exactly similar as described in a previous survey.[12] As previous survey, the interviewer-administered questionnaire covered different domains with mental health assessment being done for the first time along with STEPS surveys in India: demographic information, tobacco use was assessed using Global Adult Tobacco Survey questionnaire [13], alcohol consumption, diet and dietary salt, physical activity using Global Physical Activity Questionnaire [14], health screening, history of hypertension, diabetes, cardiovascular diseases and chronic kidney diseases, depression and suicidal behaviour, family history, health care utilization and health care costs. Assessment of depression was done by using patient health questionnaire (PHQ-9) [15] and suicidal behaviour was assessed by using the WHO module. [16]

### Physical Measurement (STEP 2)

Physiological measurements included the measurements mentioned in the above section. Blood pressure was measured using calibrated electronic equipment (OMRON HEM 7120). [17] The average of last two measurements made at intervals of two minutes, was used for analysis. The anthropometric measurements were taken using portable stadiometer and digital weighing scale matching standardised specifications recommended by WHO while undertaking such surveys (SECA, Hamburg, Germany). Weight and height of participants were determined in light clothing and without shoes. were used. Body Mass index was calculated as weight in kilograms/height in metres squared. Waist circumference (WC) and Hip circumference (HC) were measured using constant tension tape (SECA, 203). [18] WC was measured at the end of a normal expiration, with arms relaxed at the sides, at the midpoint between the lower part of the lowest rib and the highest point of the hip on the mid-axillary line.[19] Hip circumference was measured at the maximum curvature of buttocks. [19]

### Biochemical Measurement (STEP 3)

Written instructions regarding fasting, appointment date for blood test was given to the participant if selected and agreed for STEP 3. The blood samples were drawn by trained phlebotomists having a graduate or a postgraduate degree in Medical Laboratory. Blood glucose was measured using finger prick blood samples and blood glucose measurement device (Optium Freestyle).[20] Collected blood samples were centrifuged using a mini-centrifuge and separated serum was stored in ice boxes. Collected samples were transferred daily to a nearest public health institute with facility for -20°C storage. Samples were transported to the central laboratory at PGIMER, Chandigarh for analysis.

### Ethical Consideration

Ethical approval of the study was obtained from the Institute Ethics Committee of PGIMER, Chandigarh. Also, the Technical Advisory Committee of the survey approved the study protocol and also supervised the implementation and execution of the survey. Informed and written consent was taken from all the participants in the survey. Complete privacy and confidentiality of participants was assured.

### Definitions used

The cut-off criteria followed in the survey has been given in Supplementary file 1.

### Statistical analysis

Weighted analysis was conducted to calculate prevalence of NCD risk factors. Appropriate weights i.e. sampling weights, population and non-response weights were used for all data analysis to produce unbiased estimates owing to the unequal distribution of population in different strata. Separate weights were calculated for step 1 and step 3. Data cleaning as well as data analysis was done using Epi info version 3.5.2.[21] The distribution of the various risk factors were summarized as mean (SD) and frequency (proportion) depending on the type of variable. All estimates are presented with 95% confidence intervals (CIs), significance of difference in results between different groups was observed by comparing CIs. Prevalence estimates and 95% CIs were calculated using Taylor series linearization. [22]Further, data was analysed by age group, gender, and residence. Prevalence of different risk factors and proportion above standard WHO cut-off levels was determined.(Supplementary File 1). Odds ratios was calculated using multiple logistic regressions as a measure to quantify the relationship between key NCD risk factors and social determinants. SPSS version 21 [23] was used as the statistical software for analysis to accommodate for the complex survey sample design.

## Results

The response rate for STEP 1/2 and STEP 3 of the survey was 97% and 94% respectively. Out of 5250, 5115 households’ responded (99%) and 5078 individuals agreed and gave consent for STEP 1 and 2. Similarly for STEP 3, out of 2694 households 2628 responded (96%) and 2524 individuals gave consent to serum and urine sampling.

The study sample consisted of 5078 respondents, 2784 (55%) were females and 2294 (45%) were males. 68% of the study sample belonged to 18-44 years age group. Urban rural distribution of study participants is exactly similar to urban rural proportion of state population, as per Census 2011. 12% of the study participants had no formal schooling. Only 3% participants had completed post-graduation. 45% of the participants were home makers followed by non-government employees (976, 20%) and self-employed (as agriculture is the main occupation in Haryana) (857, 17%). Students, retired and unemployed (able to work) constituted 9% of the sample. The estimates of study sample characteristics are presented in <u>Table 1</u>.

**Table 1.** Socio-demographic profile of study participants in STEPS Survey, Haryana, India*

|  |  | Male (N = 2294) | Females (N = 2784) | Both Sexes (N = 5078) |
| --- | --- | --- | --- | --- |
|  |  | n (%) | n (%) | n (%) |
| Age | 18–44 | 1578(69) | 1895(68) | 3473(68) |
|  | 45–69 | 716(31) | 889(32) | 1605(32) |
| Residence | Rural | 1509(66) | 1859(67) | 3368(66) |
|  | Urban | 785(34) | 925(33) | 1710(34) |
| Education | No formal schooling | 120(6) | 394(18) | 514(12) |
|  | Less than primary school | 143(7) | 178(8) | 321(8) |
|  | Primary school completed | 326(16) | 417(19) | 743(17) |
|  | Secondary school completed | 504(24) | 374(17) | 878(21) |
|  | High school completed | 600(29) | 513(23) | 1113(26) |
|  | College/University completed | 330(16) | 245(11) | 575(13) |
|  | Post graduate degree | 58(3) | 79(4) | 137(3) |
| Social Group | SC | 807(35) | 935(34) | 1742(34) |
|  | OBC/others | 521(23) | 702(25) | 1223(24) |
|  | General | 957(42) | 1125(40) | 2082(41) |
|  | Refused | 9(0) | 22(1) | 31(1) |
| Marital Status | Never married | 392(17) | 220(8) | 612(12) |
|  | Currently married | 1852(81) | 2280(82) | 4132(81) |
|  | Separated | 6(0) | 2(0) | 8(0) |
|  | Divorced | 2(0) | 3(0) | 5(0) |
|  | Widowed | 16(0) | 255(9) | 271(5) |
|  | Refused | 26(0) | 24(1) | 50(1) |
| Occupation | Government employee | 93(4) | 45(2) | 138(3) |
|  | Non-government employee | 753(34) | 223(8) | 976(20) |
|  | Self-employed | 720(32) | 137(5) | 857(17) |
|  | Student | 156(7) | 135(5) | 291(6) |
|  | Homemaker | 81(4) | 2158(78) | 2239(45) |
|  | Retired | 37(2) | 20(1) | 57(1) |
|  | Unemployed (able to work) | 79(4) | 12(0) | 91(2) |
|  | Unemployed (unable to work) | 327(15) | 22(1) | 349(7) |
| Total |  | 2294(45) | 2784(55) | 5078(100) |

\*Figures in parenthesis indicate percentages, Abbreviations: SC: Scheduled Caste, OBC: Other Backward Castes

Mean age at initiation of tobacco smoking was 21 years and mean duration of smoking was 20 years. Prevalence of current tobacco users (smoking as well as smokeless form) was 27.4% (95% CI: 22.7-28.1). Overall, use of tobacco in the form of smoking among current users (23.5%, 21.1-25.8) of tobacco was more prevalent than smokeless form (3.9%, 2.1-4.5). (Table 2) Prevalence of smoking was high among adults aged 45-69 years old (33.1%, 28.4-37.7). More males (38.9%) smoked than females (4.3%). The prevalence of smoking was higher in rural areas (25.8%, 95% CI: 23.3-28.4) compared to those in urban areas (20.1%, 15.4-24.7). In Haryana, about 2.3 percent of the adult population was formerly smoking tobacco every day but have now stopped smoking completely. It is interesting to note that more number of current smokers who tried to quit belonged to urban areas (63%, 95% CI: 58.1-68.0), however advice by doctor was given more frequently in urban areas (33.2%, 95% CI: 22.4-43.9).

**Table 2:** Prevalence of Various NCD Risk factors in Haryana, Overall and Stratified by Age Group, Gender and Residence (Rural/Urban), 2016-2017

| Behavioural Risk Factors (%, 95% CI) |  |  |  |  |  |  |  |
| --- | --- | --- | --- | --- | --- | --- | --- |
|  | Age |  | Gender |  | Residence |  | OVERALL |
|  | 18–44 | 45–69 | Male | Female | Rural | Urban |  |
| Current tobacco Users | 20.6<br>(18.0-23.2) | 30.2*<br>(26.8-33.6) | 38.9<br>(35.3-42.4) | 4.3*<br>(3.2-5.4) | 25.8<br>(23.3-28.4) | 20.1<br>(15.4-24.7) | 23.5<br>(21.1-25.8) |
| Current drinkers | 10.6<br>(8.4-12.8) | 10.3<br>(8.5-12.2) | 18.8<br>(15.8-21.8) | 0.2*<br>(0.0-0.4) | 8.9<br>(7.4-10.5) | 12.8<br>(9.3-16.2) | 10.5<br>(8.7-12.4) |
| <5 servings of fruits and vegetables | 99.4<br>(99.0-99.8) | 99.5<br>(99.0-100.0) | 99.0<br>(98.4-99.7) | 99.5<br>(99.2-99.8) | 99.2<br>(98.9-99.6) | 99.0<br>(98.1-99.8) | 99.4<br>(99.1-99.8) |
| Low physical activity | 16.9<br>(12.6-21.3) | 13.8<br>(11.0-16.6) | 20.1<br>(15.1-25.1) | 10.9*<br>(8.2-13.7) | 15.1<br>(12.0-18.1) | 17.3<br>(9.4-25.3) | 16.0<br>(12.4-19.6) |
| Physical Measurements, (%, 95% CI) |  |  |  |  |  |  |  |
| Overweight (BMI = 25.5–29) | 23.7<br>(21.5-25.9) | 30.7<br>(25.2-36.1) | 26.8<br>(24.1-29.5) | 24.5<br>(21.8-27.1) | 22.2<br>(20.4-24.0) | 28.2<br>(26.4-30) | 25.7<br>(23.4-28.0) |
| Obesity (BMI> 30) | 7.4<br>(6.2-8.6) | 14.2*<br>(11.1-17.4) | 6.2<br>(4.6-7.8) | 13.4*<br>(11.8-15.1) | 7.3<br>(6.2-8.5) | 11.5*<br>(10.4-12.7) | 9.4<br>(8.0-10.9) |
| Abdominal Obesity (Males>90, Females>80) | M= 17.9<br>(15.3-20.4) | M= 31.4<br>(24.0-38.8) | 21.8<br>(18.2-25.1) | 45.1*<br>(41.6-48.6) | 9.49<br>(8.1-10.9) | 22.1*<br>(19.3-24.8) | NA |
|  | F= 36.7<br>(32.4-41.1) | F= 64.0<br>(59.1-68.9) |  |  |  |  |  |
| Elevated Blood Pressure (SBP>140 and/or DBP>90) | 20.7<br>(19.1-22.3) | 39.1*<br>(35.9-42.3) | 29.5<br>(26.8-32.2) | 22.1*<br>(20.0-24.2) | 24.4<br>(22.3-26.4) | 28.8<br>(26.0-31.5) | 26.2<br>(24.6-27.8) |
| Biological Risk Factors, (%, 95% CI) |  |  |  |  |  |  |  |
| Hyperglycemia (>110mg/dl) | 15.0<br>(12.1-18.0) | 21.3<br>(16.4-26.2) | 14.2<br>(9.7-18.7) | 17.6<br>(12.1-23.2) | 12.6<br>(10.0-15.2) | 19.7<br>(9.5-29.9) | 15.5<br>(11.0-20.0) |
| Hypertriglyceridemia (>150mg/dl) | 29.8<br>(25.9-33.7) | 39.5<br>(32.5-46.5) | 36.4<br>(31.9-40.8) | 25.8*<br>(22.2-29.5) | 33.0<br>(28.8-37.2) | 31.4<br>(27.1-35.6) | 32.3<br>(29.3-35.4) |
| Hypercholesterolemia (>190mg/ dl) | 27.0<br>(23.2-30.9) | 43.9*<br>(36.4-51.4) | 30.6<br>(25.9-35.2) | 33.3<br>(29.1-37.5) | 30.4<br>(26.1-34.6) | 33.5<br>(27.0-40.0) | 31.6<br>(28.0-35.3) |
| Raised salt intake (>5gm/day) | 91.3<br>(89.1-93.5) | 91.5<br>(88.7-94.3) | 94.5<br>(92.6-96.3) | 86.4*<br>(83.1-89.4) | 89.3<br>(87.4-91.2) | 94.4<br>(90.7-98.1) | 91.3<br>(89.3-93.3) |

Prevalence of current alcohol use in the state was 10.5% (95% CI: 8.7-12.4) without any rural urban differences. (Table 2). Among males, 18.8% (15.8-21.8) were current users. 3.1% among males and none of the females consumed higher levels of alcohol. Harmful alcohol consumption was found in 0.1% (0.0-0.2) of the population. Overall no significant differences could be found in alcohol consumption patterns in urban and rural areas.

Lower levels of intake of fruits and vegetables were found to be high among both age groups, both sexes as well as residence. Overall 99.2 % (95% CI: 98.9-99.6) of participants took less than 5 servings of fruits and/or vegetables on average per day. In a typical week, fruits and vegetables are consumed on 1 and 4 days respectively. 7.2% of the population (95% CI: 6.0-8.4) always/often added salt before/when eating (rural significantly more than urban).

Low levels of physical activity, i.e. activity levels of less than 600 MET minutes were prevalent among 11% (95% CI: 8.6-13.4) of the respondents. Compared to females (6.0%, 4.2-7.8), males had a higher prevalence of low physical activity (15.1%, 11.5-18.7). Rural (9.5%) as well as urban (13.1%) areas had people with low physical activity.(<u>Table 2</u>)

Overweight and obesity combined was observed in 35.2% (95% CI: 32.6-37.7) of participants. 45-69 years age group had significantly higher number of people having overweight (30.7%, 25.2-36.1) as well as obesity (14.2%, 11.1-17.4). The central obesity was found to be higher among females (73.8%, 70.3-77.3) than males (53.3%, (95% CI: 49.7-56.9).

The prevalence of elevated blood pressure (including those who were on medication for hypertension) was 26.2% (95% CI: 24.6-27.8). Significantly higher prevalence was observed among those aged 45–69 years (39.1%, 35.9-42.3), males (29.5%, 26.8-32.2), as compared to those aged 18-44 years (20.7%, 19.1-22.3) and females (22.1, 95% CI: 20.0-24.2).

It was found that 15.5% (95% CI: 11.0-20.0) of the study participants had raised blood glucose. The prevalence was higher among those aged 45–69 years (22.7%, 18.1-27.2). No difference was found in prevalence by gender and residence.

Hypercholesterolemia and Hypertriglyceridemia was found to be in 31.6% (95% CI: 28.0-35.3) and 32.3% (95% CI: 29.3-35.4) of population respectively. For both the parameters, values were higher for 45-69 years old, males and rural populations, though none of the parameters had significant differences.

The salt intake was found to be higher i.e. more than 5 grams per day for 91.3% (95% CI: 89.3-93.3) of population. It is interesting to note that out of 46 % who said they feel they consume just the right amount of salt had daily intake of salt more than 5gm/day.

The mean values of different parameters for behavioral risk factors, physical measurements are presented in **Table 3**. An average of 10 (95% CI: 8.0-12.0) cigarettes were being consumed per day by a daily smoker. Similarly average number of standard drinks being consumed by current drinkers were 3.2 (95% CI: 3.0-3.5). The mean value of BMI was 24.6 (95% CI: 23.3-24.9) and mean differed significantly between males and females. There was no difference in mean waist hip ratio by age, gender and residence.

**Table 3:** Means (Confidence intervals) of different parameters for behavioral information and physical measurements in population of Haryana, 2016-2017

| Mean of different behavioural risk factors on average per day, Mean (95% CI) |  |  |  |  |  |  |  |
| --- | --- | --- | --- | --- | --- | --- | --- |
|  | Age |  | Gender |  | Residence |  | OVERALL |
|  | 18–44 | 45–69 | Male | Female | Rural | Urban |  |
| Mean amount of tobacco used by daily smokers on average per day | 9.8<br>(7.2-12.4) | 10.6 | 10.2 | 4.9*<br>(2.3–7.4) | 10.6 | 9.2 | 10.0<br>(8.0–12.0) |
|  |  | (8.7-12.5) | (8.2-12.3) |  | (9.1-12.1) | (4.7-13.6) |  |
| Mean number of standard drinks consumed on average per day | 3.1<br>(2.7-3.4) | 3.6<br>(3.1-4.1) | 3.3<br>(3.0-3.5) | 2.4<br>(1.8-3.0) | 3.4<br>(3.0-3.7) | 3.1<br>(2.7-3.5) | 3.2<br>(3.0-3.5) |
| Mean number of servings of fruits and/or vegetable on average per day | 1.5<br>(1.5-1.6) | 1.5<br>(1.4-1.5) | 1.5<br>(1.5-1.6) | 1.4<br>(1.4-1.5) | 1.4<br>(1.4-1.5) | 1.6<br>(1.5-1.7) | 1.5<br>(1.4-1.5) |
| Mean number of servings of fruits on average per day | 0.6<br>(0.6-0.7) | 0.6<br>(0.5-0.6) | 0.6<br>(0.6-0.7) | 0.6<br>(0.5-0.6) | 0.5<br>(0.5-0.6) | 0.7<br>(0.6-0.8) | 0.6<br>(0.5-0.6) |
| Mean minutes spent in sedentary activities on average per day | 53.0<br>(50.7-55.2) | 53.2<br>(50.1-56.3) | 52.5<br>(49.2-55.9) | 53.7<br>(50.8-56.6) | 52.6<br>(50.4-54.8) | 53.6<br>(50.7-56.4) | 53.0<br>(51.3-54.8) |
| Mean minutes of total physical activity on average | 217.5<br>(205.8-229.2) | 216.6<br>(202.1-231.0) | 179.7<br>(167.8-191.6) | 263.8*<br>(248.9-278.7) | 239.8<br>(227.9-251.7) | 185.5*<br>(168.1-203.0) | 217.2<br>(206.2-228.2) |
| Mean of different physical parameters, Mean (95% CI) |  |  |  |  |  |  |  |
| Body Mass Index (BMI) | 23.1<br>(22.9-23.4) | 24.6*<br>(24.1-25.1) | 23.4<br>(23.0-23.8) | 24.2<br>(23.8-24.5) | 23.1<br>(22.8-23.3) | 24.6*<br>(24.0-25.1) | 24.6<br>(23.3-24.9) |
| Waist Circumference (WC) | 79.6<br>(78.7-80.5) | 84.8*<br>(83.9-85.7) | 82.6<br>(81.9-83.4) | 78.8*<br>(78.1-79.6) | 79.8<br>(79.1-80.4) | 82.6*<br>(81.7-83.6) | 81.2<br>(80.3-82.1) |
| Hip Circumference (HC) | 90.9<br>(89.8-92.1) | 94.6*<br>(93.4-95.7) | 91.5<br>(90.8-92.2) | 92.1<br>(91.4-92.8) | 90.5<br>(89.9-91.1) | 93.5*<br>(92.6-94.4) | 92.0<br>(90.7-93.4) |
| Waist Hip Ratio (WHR) | 0.9<br>(0.9-0.9) | 0.9<br>(0.9-0.9) | 0.9<br>(0.8-0.9) | 0.9<br>(0.9-0.9) | 0.9<br>(0.9-0.9) | 0.9<br>(0.9-0.9) | 0.9<br>(0.9-0.9) |
| Systolic Blood Pressure (SBP) | 120.7<br>(120.0-121.4) | 130.8*<br>(129.3-132.3) | 126.3<br>(125.1-127.4) | 120.5*<br>(119.7-121.4) | 123.1<br>(122.3-123.9) | 124.6<br>(123.3-125.9) | 123.7<br>(123.0-124.4) |
| Diastolic Blood Pressure (DBP) | 81.2<br>(80.6-81.7) | 86.0*<br>(85.3-86.7) | 83.7<br>(82.9-84.5) | 81.2*<br>(80.6-81.8) | 82.1<br>(81.5-82.7) | 83.3<br>(82.5-84.2) | 82.6<br>(82.1-83.1) |
| Mean of different biological parameters, Mean (95% CI) |  |  |  |  |  |  |  |
| Glucose | 95.0<br>(92.6-97.5) | 104.2*<br>(100.9-107.5) | 96.8<br>(94.4-99.1) | 98.8<br>(95.9-101.6) | 95.2<br>(93.1-97.4) | 100.8<br>(96.9-104.7) | 97.5<br>(95.4-99.7) |
| Triglycerides | 135.7<br>(127.3-144.1) | 148.0<br>(135.6-160.4) | 145.7<br>(135.1-156.2) | 128.0<br>(121.4-134.5) | 140.2<br>(131.3-149.1) | 137.1<br>(125.5-148.7) | 138.9<br>(131.8-146.0) |
| Cholesterol | 170.4<br>(165.2-175.7) | 190.3*<br>(181.6-198.9) | 173.9<br>(167.9-179.9) | 178.9<br>(173.0-184.8) | 175.6<br>(171.4-179.8) | 176.1<br>(164.2-188.0) | 175.8<br>(170.5-181.1) |
| Salt intake | 8.1<br>(7.8-8.3) | 8.0<br>(7.7-8.3) | 8.8<br>(8.6-9.0) | 6.9<br>(6.7-7.1) | 7.8<br>(7.6-7.9) | 8.5<br>(8.1-8.8) | 8.0<br>(7.9-8.2) |

It is important to note that 13.1%, 4.3% and 1.8% of respondents were known hypertensive, diabetic or hyperlipidaemia patients, (Table 4) and only 58%, 84% and 1.5% were on medicines for above mentioned diseases respectively. 3.1%, 5.6% and 3.1 % of respondents had visited traditional healers for their ailment respectively.

**Table 4.** Self-reported prevalence of NCDs in Haryana, 2016-2018

| Known Chronic Disease History (% diagnosed in past 12 months or earlier) |  |  |  |  |  |  |  |
| --- | --- | --- | --- | --- | --- | --- | --- |
|  | Age |  | Gender |  | Residence |  |  |
|  | 18-44 | 45-69 | Male | Female | Rural | Urban | Overall |
| Hypertension | 8.7<br>(7.2-10.2) | 23.4*<br>(19.3-27.5) | 9.7<br>(7.7-11.7) | 17.2*<br>(15.0-19.4) | 11.4<br>(9.6-13.2) | 15.4<br>(12.2-18.7) | 13.1<br>(11.4-14.8) |
| Diabetes | 1.5<br>(1.0-2.1) | 10.9*<br>(6.8-14.9) | 4.5<br>(2.9-6.2) | 4.0<br>(2.5-5.6) | 2.8<br>(2.1-3.5) | 6.4<br>(4.2-8.7) | 4.3<br>(3.1-5.5) |
| Hyperlipidemia | 1.1<br>(0.6-1.5) | 3.6<br>(1.4-5.9) | 1.8<br>(1.0-2.7) | 1.8<br>(1.0-2.6) | 1.2<br>(0.7-1.8) | 2.7<br>(0.9-4.5) | 1.8<br>(1.0-2.6) |
| Depression* | 1.5<br>(0.8-1.8) | 1.3<br>(0.6-1.9) | 1.1<br>(0.6-1.6) | 1.6<br>(1.0-2.2) | 1.0<br>(0.8-1.8) | 1.8<br>(0.5-2.1) | 1.4<br>(0.9-1.8) |

\*Severe depression symptoms, person scoring 18-27 in the PHQ9 module

About one-third (34.5%) of the participants had mild to moderate levels of depression and 1.4 % of the participants were found to have severe depression, needing immediate medical attention Additionally, 5% respondents agreed attempting suicide in the last 12 months. A higher percentage of respondents in 18-44 years age group and those residing in rural areas had considered though differences were not statistically significant. Only 14% among these respondents sought professional help.

It is interesting to note that though known hypertensive, diabetic or CVD patient are prevalent in rural as well as urban areas, differences in prevalence by residence is significantly higher for hypertension, CVD and diabetes. Prevalence is higher for urban areas than rural areas except for CVD.

A significant number of subjects reported a family history of hypertension (42.6%), diabetes (20.8%), chronic kidney diseases (4.7%), early myocardial infarction (3.2 %), cancer or malignant tumor (3.7%), and raised cholesterol (3.6%), with 30 % of participants reporting any of these NCDs.

Another important aspect of awareness levels as well as unmet need on part of population is the fact that despite such high prevalence of hypertension, 43.8% of the population had never got their blood pressure measured. Also, of all the known hypertensive cases in the state, 33.4% of the respondents were aware of their condition, 26.3% are on treatment while only 12% of the cases are controlled. (**Error! Reference source not found.**) For diabetes the condition is even more serious, of all the known diabetes cases in Haryana 29.5% of the respondents were aware of their condition, 22.4% are on treatment while only 13.8% of the cases of Diabetes are controlled.

**Fig 1.**
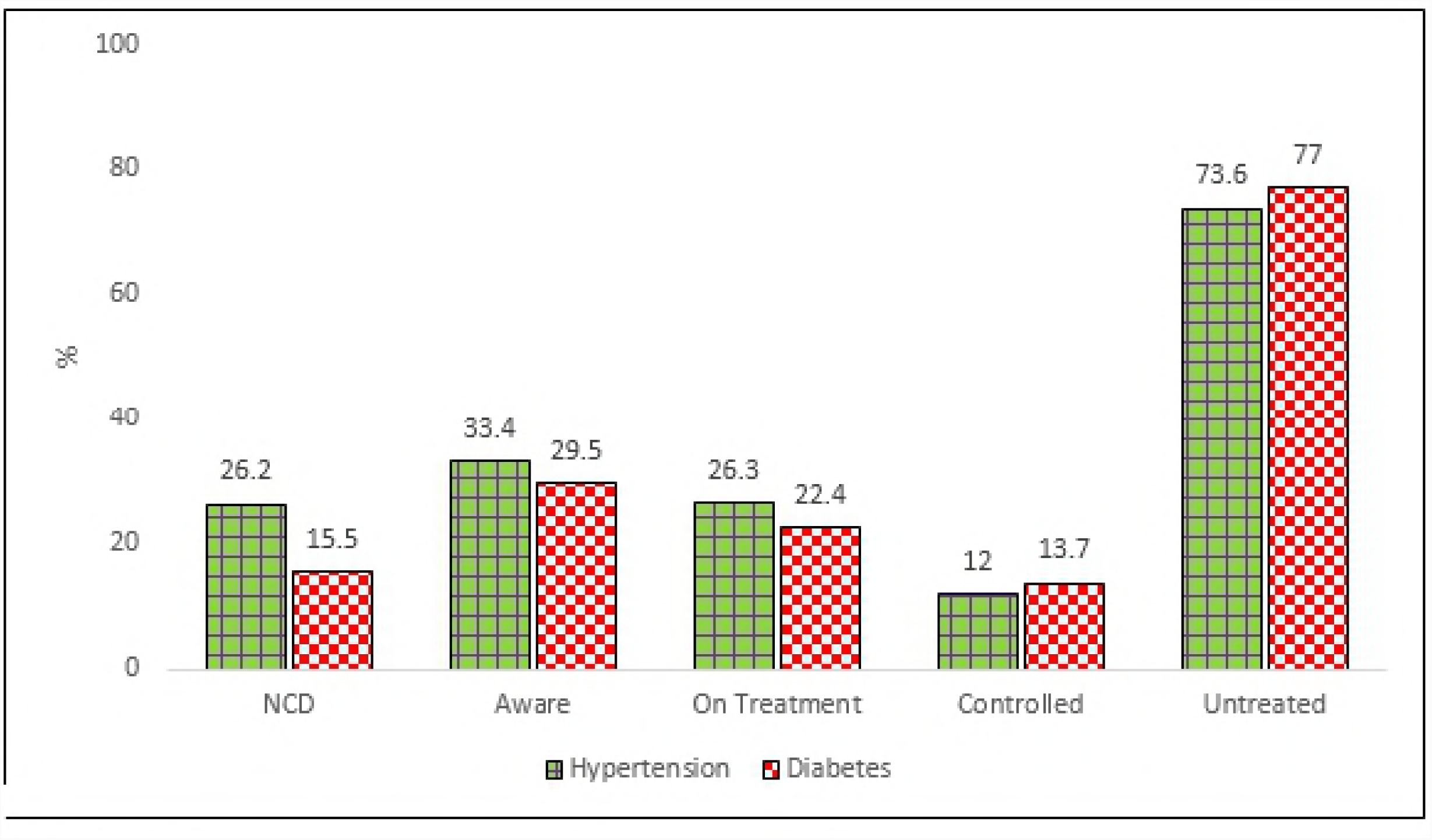
Status of Hypertension and Diabetes: Extent of awareness, treatment and controlled cases in Haryana, 2016-2018

On asking regarding utilization of preventive services for different cancers, only 7.7% and 8% of females aged 30-49 were ever screened for cervical or breast cancer. 14.7% (95% CI: 12.0-17.5) respondents reported to have been screened for oral cancer.

Only 0.3% (95% CI: 0.1-0.4) of the population was completely free from the 5 established risk factors. More than three fourth of the respondents (76.4 %) had at least 1-2 risk factors and one third (23.4%) had 3-5 risk factors.

Mean amount of INR 2096 and 6794 is spent by respondents for OPD consultations (last 30 days) or hospitalizations (last 1 year) due to NCDs average INR 1604 is spent for health care availed not related to any visits/hospitalization.

Owing to limited scope of describing all the results in a single manuscript, the health care costs, mental health, and other domains will be described in subsequent publications.

Results of multiple logistic regression analyses is shown in <u>Table 5</u> highlight the increasing prevalence of different risk factors with age, adjusting for other factors including gender and the place of residence. Males were found to be having higher odds of being a current smoker and drinker while females were found to be having about twice the odds of obesity and low physical activity. While the urban residence was associated with higher odds of obesity, hyperglycaemia and hypercholesterolemia

**Table 5:** Determinants of NCD risk factors among aged 18–69 years in the State of Haryana, 2016-2018

|  |  | Current<br>Smokers<br>OR<br>(95% CI) | Current<br>Drinkers<br>OR<br>(95% CI) | Low Physical<br>Activity OR<br>(95% CI) | Insufficient<br>Servings<br>OR<br>(95% CI) | Raised Blood<br>Pressure OR<br>(95% CI) | Obesity<br>OR<br>(95% CI) | Raised<br>Glucose<br>OR<br>(95% CI) | Raised<br>Cholesterol<br>OR<br>(95% CI) |
| --- | --- | --- | --- | --- | --- | --- | --- | --- | --- |
| Education | No formal<br>Schooling | 1 | 1 | 1 | 1 | 1 | 1 | 1 | 1 |
|  | Primary<br>Education | 0.3<br>(0.2-0.5) | 1.2*<br>(0.8-1.8) | 0.3(0.1-1.2) | 1.5*(1.2-1.9) | 1.6*(1.3-2.1) | 1.0(0.7-1.4) | 1.4*(1.0-1.9) | 1.5(0.7-3.2) |
|  | Secondary<br>Education | 0.5<br>(0.4-0.7) | 0.9*<br>(0.7-1.6) | 0.6(0.3-1.3) | 1.2*(1.0-1.5) | 1.3*(1.1-1.6) | 1.0(0.7-1.4) | 0.9(0.7-1.10) | 0.9(0.5-1.70) |
|  | Higher<br>Education | 0.9*<br>(0.7-1.1) | 1.2*<br>(0.7-1.5) | 1.2(0.7-2.0) | 1.3*(1.0-1.5) | 1.5*(1.2-1.8) | 0.8(0.6-1.0) | 1.1(0.9-1.4) | 0.7(0.4-1.3) |
| Social<br>Group | General | 1 | 1 | 1 | 1 | 1 | 1 | 1 | 1 |
|  | Scheduled<br>Caste | 1.1*<br>(0.9-1.4) | 0.9*<br>(0.7-1.1) | 1.3(0.7-2.3) | 1.2*(1.0-1.4) | 1.4*(1.2-1.7) | 1.8*(1.3-2.5) | 1.6*(1.3-2.0) | 1.2(0.7-2.1) |
|  | Other<br>Backward | 0.7<br>(0.5-0.9) | 0.6<br>(0.5-0.8) | 0.9(0.5-1.8) | 1.3*(1.1-1.6) | 1.4*(1.2-1.7) | 1.7*(1.2-2.3) | 1.4*(1.1-1.8) | 1.1(0.6-1.9) |
| Gender | Males | 1 | 1 | 1 | 1 | 1 | 1 | 1 | 1 |
|  | Females | 0.07<br>(0.05-0.1) | 0.04<br>(0.02-0.07) | 1.3 (0.6-2.9) | 0.8*(0.6-1.0) | 1.1(0.8-1.4) | 2.2*(1.4-3.3) | 0.9(0.7-1.2) | 0.8(0.6-1.1) |
| Age group | 18–44 | 1 | 1 | 1 | 1 | 1 | 1 | 1 | 1 |
|  | 45–69 | 1.8<br>(1.5-2.2) | 1.2*<br>(1.0-1.5) | 1.1(0.6-2.1) | 1.2*(1.0-1.5) | 0.9(0.7-1.0) | 1.9*(1.4-2.4) | 0.9(0.8-1.2) | 1.7*(1.0-2.8) |
| Marital Status | Currently married | 1 | 1 | 1 | 1 | 1 | 1 | 1 | 1 |
|  | Never married | 1.8<br>(1.1-3) | 0.5*<br>(0.2-1.3) | 0.8(0.2-3.5) | 1.0(0.7-1.4) | 1.1(0.7-1.5) | 1.1(0.7-1.8) | 1.0(0.6-1.5) | 1.2(0.5-3.1) |
|  | Divorced/<br>Widowed | 3.3<br>(1.9-6) | 1.2*<br>(0.5-3.0) | 1.1(0.2-5.6) | 1.1(0.7-1.7) | 0.9(0.6-1.4) | 2.4(1.1-5.1) | 0.9(0.5-1.6) | 1.6*(0.4-5.5) |
| Residence | Rural | 1 | 1 | 1 | 1 | 1 | 1 | 1 | 1 |
|  | Urban | 0.6<br>(0.5-0.7) | 1.0*<br>(0.8-1.2) | 0.7(0.4-1.10) | 1.0(0.9-1.2) | 1.1(1.0-1.3) | 1.9*(1.5-2.5) | 1.4*(1.2-1.7) | 2.1*(1.3-3.2) |
| Occupation | Home Maker | 1 | 1 | 1 | 1 | 1 | 1 | 1 | 1 |
|  | Government Employee | 2.4<br>(1.6-3.7) | 3.6<br>(2.1-6.4) | 0.3*(0.1-0.8) | 1.3(1.0-1.7) | 1.0(0.8-1.4) | 0.890.5-1.40 | 1.0(0.7-1.5) | 0.9(0.3-2.3) |
|  | Non-Govt. Employee | 1.7<br>(1.0-2.7) | 1.0*<br>(0.6-1.5) | 0.8(0.2-3.6) | 1.0(0.6-1.7) | 1.0(0.6-1.7) | 1.0(0.4-2.7) | 1.5(0.7-3.2) | 0.4(0.1-1.3) |
|  | Self employed | 1.0*<br>(0.8-1.3) | 1.0*<br>(0.8-1.3) | 0.6(0.3-1.2) | 1.1(0.9-1.5) | 1.0(0.8-1.3) | 1.2(0.7-2.2) | 1.2(0.9-1.7) | 0.9(0.4-2.4) |
|  | Others | 1.2*<br>(1.0-1.6) | 1.0*<br>(0.8-1.2) | 0.7(0.4-1.5) | 1.2(0.9-1.5) | 1.0(0.8-1.3) | 0.6(0.4-1.0) | 0.9(0.6-1.2) | 0.7(0.3-1.60) |

## Discussion

This state wide community based survey has demonstrated high rates for several NCD risk factors in adult population in Haryana, India. It is one of the first STEPS survey in the world using m STEPS android application. [24] The current study has generated data on prevalence of different risk factors prevalence by age group, gender and place of residence which also allowed to explore their relation with socio demographic indicators. The methodology adopted for this survey is in accordance with international recommendations and allows comparison with similar STEPS surveys [25-28] as in the previous survey in neighbouring Indian states. [12, 29, 30]

The prevalence of current tobacco smokers (23.5%) and smokeless tobacco users (3.9%) was consistent with the estimates of GATS for the state i.e. 19.7% and 6.3% respectively.[6] However, the smokeless tobacco rates in the state are much lower than the national average of 21.4% and several other states primarily the North East Indian states owing to their cultural practices of consuming betel nut called *tamol.*[31] However, the prevalence of current tobacco smoking is high in comparison to several neighbouring states as well as national average of India (10.7%).[6] Contrary to common notion against use of tobacco by females in India, females in Haryana were found to be consuming tobacco at rates twice more (4.3%) than national average (2%). Socio-cultural influences may be the possible attribute to this pattern. Our findings of higher proportion of quitters in urban areas are consistent with recent literature from similar settings [32]. Consistent with previously reported literature, the plausible explanation for this could be better availability and accessibility of tobacco cessation measures in the state.[33] Efforts to control tobacco consumption have been initiated in the state [34], however, demand further strengthening of the policies as well as their ground level implementation. This is especially in light of similar prevalence rates in few other lower-middle income countries in Asian continent who have failed to curb the tobacco menace despite decades of consistent efforts.[35]

The current study also revealed lower percentage of alcohol users (10.5 %) than neighbouring North Indian state.[12] Of these current drinkers, with 1.3% of them being high-end users (≥ 60 g). Prevalence of alcohol consumption among males (18.8%) in survey population was slightly lower than to those of a NFHS 2015-16 in which 24% males were found to be current drinkers.[36-38] Harmful use of alcohol, though prevalent in only about 3% of the total study population, is more than twice as high among those aged 45-69 years, compared to those aged 18-44 years; a finding that is in agreement to what has been observed in the other surveys and is in contrast to the belief that alcohol use is a problem among reckless youth and prevalence declines as people mature and take responsibilities.[39] In Haryana, 79% of the total respondents were lifetime abstainers in line with the global trends that show particularly high levels of alcohol abstinence especially across North Africa and the Middle East.[40] Social drinking is perceived to reduce stress and anxiety, however, studies reveal that in addition to being a risk factor for various NCDs, alcohol consumption contributes to other class of mortality burden including road crashes and injuries.[40] Patterns of alcohol use identified in the survey are detrimental to health of people of Haryana and need corrective measures. [41]

Global Burden of Disease Study also identified dietary risk factors as third common risk factor in Haryana for morbidity and mortality.[6] The current survey second this finding as it was found that about 99% of the study population consumed less than the recommended 5 servings daily of fruits and vegetables. The low intake of fruits and vegetables has been reported in various studies in India and also globally.[29, 42-44] Though prevalence of low fruit and vegetable intake may be considered an availability and accessibility issue (urban area residents consume more fruits and vegetables) but this difference could be by choice as Haryana is the top consumer of dairy and its products in the country (thrice than the national average).[45]

The current study findings on lower percentage of those not meeting WHO recommendations (11%) points towards higher physical activity levels in comparison to global levels of physical activity.[44] These levels were particularly work related activities and that also among females than males. The fact that levels were found to be higher than neighbouring states point towards different socio cultural patterns where the females in rural areas are still relying on non-technological mechanisms. Since Haryana is predominantly an agriculture driven state with several belts still under development stages,[46] low physical activity levels are not as high as in the other states of the country.[12, 30]

Despite good physical activity levels, high prevalence of being overweight (26%) and obesity (9%), which is higher among females (24.5% overweight, 13.4% obese) than males (26.8% overweight, 6.2% obese) points towards detailed exploration of other behavioural risk factors such as dietary fat intake. The higher prevalence is consistent with results of worldwide prevalence of obesity and is twice as compared to national figures (4.9%).[47] That one in every three persons in Haryana is overweight is an indication towards the need for reviewing current policies in school, work places and other targeted setting. If Asian cut-offs [48] for obesity are followed, the prevalence of obesity (21.7%) and overweight (31.2%) is comparable to obesity levels in many developed countries. [49-52]

High prevalence of hypertension (26%), hypercholesterolemia (31.6%) hypertriglyceridemia (32.3%) is alarming and underscores that these are key risk factors for NCDs. In Global Burden of Disease study (Haryana estimates), these have been identified as one of the top leading risk factors for NCDs burden in the state, though their figures are lower than survey findings and ranks are in reverse.[6] It has to be understood that the GBD estimates relied upon several small non representative datasets across Haryana and there is need for more robust estimates as generated under this survey.

In the current study, tobacco smoking, alcohol drinking and raised blood pressure (BP) were more prevalent in males than females which is consistent with findings from several other STEPS surveys in the country as well as surrounding nations.[27, 43, 53, 54] Despite higher physical activity levels among females, the central as well as abdominal obesity were higher among females. This calls for an in-depth understanding of dietary patterns and type of physical activities among females. The mean number of days and servings of fruit consumed, the physiological measurements including the BMI, waist circumference and percentage of population who are overweight and obese are significantly higher in the urban areas than those in the rural areas. The percentage of people who consume tobacco daily and add more salt to their food before eating, were found to be significantly higher in rural areas than the urban areas.

Another important aspect of awareness levels as well as unmet need on part of population was the fact that despite such high prevalence of hypertension, 43.8% of the population had never got their blood pressure measured. It is worth noticing that of all the known hypertensive cases, 33.4% were aware of their condition, 26.3% were on treatment while only 12% of the cases were controlled. So the screening process which has already been undertaken in the state needs to be complemented and supplemented with awareness campaigns and treatment adherence policies or follow-up strengthening.[55]

The observed prevalence of diabetes in the current study was 15.5%, which is higher than estimates for global (9%) as well as South Asia overall (8%).[56, 57] However, prevalence has been observed to be present uniformly across both genders, age groups and residence status of population. NFHS-4, which is a national level survey in the country for the first time reported the prevalence of diabetes which are much lower and can be attributed to its higher cut-off criteria to determine the raised blood glucose levels, different age-group of the sampled population and also due to the calculation of random blood glucose as against the standard practice of calculating fasting blood glucose.[38]

Assessment of mental health status undertaken in this survey revealed the need for mental health interventions as despite 5% of the population having moderate to severe depression, none of them reported any consultation/ expenditure on this group of diseases. It is cardinal to focus on this under-represented aspect of association between mental health and other noncommunicable diseases such as cardiovascular diseases and obesity and hence increased risk of mortality as reported in various studies. [58]. The National Mental Health Survey 2015-16 reported that nearly 11% of adult Indians suffer from some form of mental disorders and most of them do not receive care for a variety of reasons.[59] Mental health is a neglected area in developing countries, despite on-going programmes. Our survey was a first attempt to demonstrate feasibility of including mental health for a sub population as a part of STEPS surveys. The National Mental Health Survey 2015-16 provided estimates of only 12 states of the country in which Haryana was not included in the sampling frame. [60]

Interventions involving direct family members of a person having NCD will be required as risk factors for this group of disease are mostly a result of family choices and not individual. 40-50% of those suffering from any of the common NCDs reported having family members with hypertension. This was closely followed by diabetes (25-35%). In addition, 3% of the population had a family member who has suffered early myocardial infarction (early myocardial infarction < 55 years). This points towards emergence of NCDs in younger age groups.

Uptake or availability of screening services for different cancers is higher than Punjab.[12] A probable reason could be maturity of NPCDCS program in Haryana since its launch, as services got strengthened in 2017 as compared to 2015 when Punjab survey was conducted. Further explorative research needs to be undertaken to attribute the current levels of screening to the implementation of program.

The fact that only 0.2% of the study population was found to be free of all studied NCD risk factors is an indication of growing epidemic of NCDs in the state. The results are in line with other studies where less than 1% of the population is free from any of the risk factors for NCDs. [25, 42, 61]. The maximum proportion of the population have at least 1-2 risk factors. Proportion of 40–69 year old adults with a 10-year risk of cardiovascular disease ≥30% was also substantial at 17.7 %, with the proportion almost double (33.4%) among the 55–69 year old age group as compared to 7% in a neighbouring state. High consumption of tobacco contributes towards this raised risk.[62]

Global burden of diseases study has documented high levels of epidemiologic transition in different nations with huge variation among Indian sub-populations.[6] In line with this epidemiological transition, the composition of risk factors that drives its disease burden has also changed over time. The GBD estimates of NCD risk factors and that of community based surveys in different states differ a lot. Many states which started implementation of program fail to document effectiveness of their interventions due to lack of baseline levels of NCDs in their populations.

### Strengths and Limitations

We have successfully demonstrated the use of additional modules of CKD, mental health assessment using PHQ-9, [15] health care costs for all NCDs. Additional questions on COPD/asthma or NCDs in females or geriatric population could be added with minimal change in the existing format as in the LMICs chronic respiratory diseases are a major problem. Owing to the resource constraints, the study was limited to 18-69 years age group, however, many of the behavioral risk factors such as tobacco and alcohol. Also, there could be a possibility to design a mechanism for telephonic interviews for the participants not available for an interview throughout the day or subsequent days. The possibility of entering the data into the application by the participants themselves could be explored.

## Conclusion

NCD risk factors are uniformly prevalent in the population of Haryana. The estimates which have been generated in the survey will contribute towards development and evaluation of state NCD control program. The estimates generated by this survey provided baseline data for state wide action plan prepared by state for specific population and individual health interventions for implementation. It gives baseline data for planning the program and devising setting specific well-tailored interventions. The use of STEPS methodology will enable future state wide, national and international comparisons. Survey will also provide data for 12 indicators of National NCD monitoring framework out of 21 indicators which is first of its kind in India. The use of STEPS methodology will enable future state-wide, national and international comparisons. The results steers the attention towards the need for political commitment and increase in resource allocation for NCDs to successfully achieve the sustainable development goals and National Monitoring Framework targets required to monitor the NCD crisis.

